# Improved sensors for fructose-1,6-bisphosphate enable in vivo imaging of glycolysis

**DOI:** 10.64898/2026.04.29.721630

**Authors:** Juliann Tyler, Anjali Amrapali Vishwanath, Triveni Menon, Tanishqua Duarah, Raghabendra Adhikari, John N. Koberstein, Daniel Feliciano, Isabel Espinosa-Medina, Daniel Colón-Ramos, Alison G. Tebo

**Affiliations:** Howard Hughes Medical Institute – Janelia Research Campus, Ashburn, VA, USA; Department of Neuroscience and Cell Biology, Yale University School of Medicine, New Haven, CT 06536; Wu Tsai Institute, Yale University, New Haven, CT 06510

## Abstract

Fructose-1,6-bisphosphate (FBP) is the product of the first committed step of glycolysis, and its concentration is tightly correlated with glycolytic flux. Glycolytic activity varies across tissues and cell types: some tissues, such as the brain, dynamically regulate glycolysis in response to demand, while others, such as the liver have characterized spatial heterogeneity. Here, we report HYlight2, an improved sensor for FBP developed through random whole-gene mutagenesis in *E. coli* lysate. After four rounds of screening, we isolated HYlight2, which retains its binding affinity while displaying a ΔR/R ∼9 *in vitro*, a three-fold improvement in mammalian cells, and a two-fold improvement in detecting glycolytic responses during stimulated neuronal activity. We further demonstrate its use *in vivo* to detect altered glycolytic activity in *C. elegans* neurons, zebrafish pancreatic islets, and mouse liver.

## Introduction

Metabolism exhibits high spatiotemporal heterogeneity^1,2^ however, the tools to measure are lacking and often involve invasive sample preparation. In particular, mass spectrometry imaging-based measurements are pushing our understanding of metabolic heterogeneity,^3–5^ although these techniques still suffer from poor temporal resolution and struggle to resolve subcellular distributions. Furthermore, different cell types often exhibit metabolic pathway preferences. Certain cell types rely heavily on glycolysis, while others, like neurons, dynamically regulate glycolysis in response to energy demands.^6,7^ The intermediate fructose 1,6-bisphosphate (FBP), produced by phosphofructokinase (PFK), is the first committed step of glycolysis, and its intracellular concentration has been shown to correlate with flux through the glycolytic pathway.^8,9^ Thus, spatiotemporally resolved measurements of FBP provide an avenue towards understanding the heterogeneity of glycolysis and glucose utilization in different cell types, tissues, and organisms.

To this end, a genetically encoded fluorescent biosensor named HYlight was developed.^10^ HYlight was generated via the insertion of a circularly permuted green fluorescent protein (cpGFP) into the protein CggR, a bacterial transcription factor that binds FBP. This sensor offers ratiometric excitation contrast and provides a method for imaging glycolytic dynamics in cells as changes in fluorescence ratio (ΔR/R0) track corresponding changes in the concentration of FBP. While HYlight has proved useful for measurement of glycolytic dynamics in cell lines and *C. elegans*,^7^ a modest dynamic range and lack of affinity variants have impeded its widespread implementation. To address these issues, we used directed evolution to engineer HYlight2, which shows improved dynamic range while retaining its binding affinity and ratiometric excitation contrast. Furthermore, we demonstrate that HYlight2 is 2.5x more sensitive for detection of glycolytic changes resulting from stimulated neuronal activity in cultured neurons. We also benchmarked HYlight2 against HYlight to detect physiological responses from single neurons in *C. elegans*, fluctuations in FBP in developing zebrafish pancreatic islets, and metabolic zonation in mouse liver.

## Results

### Evolution of FBP Biosensor HYlight2

HYlight was originally engineered by inserting cpGFP into the binding domain CggR and extensively screening combinations of the two residue linkers flanking cpGFP (Figure 1A). We reasoned that whole gene mutagenesis would allow us to further improve the sensor by exploring sequences that were not screened in the previous library design. The number of possible protein sequences accessible via random mutagenesis is far larger than can be adequately screened, and this limits the amount of exploration of the sequence-fitness landscape that is possible. In this study, we experimented with “population splitting”,^11^ which involves dividing screening bandwidth into four smaller efforts that explore the sequence landscape independently. Thus, where one library trajectory may get stuck in a local extremum, the additional exploration of the sequence space conferred by the parallel trajectories can still allow for beneficial and improved sequences to be identified (Figure 1B). Using error prone PCR, we used a high-throughput assay to screen for FBP-responsive variants in bacterial lysate (-/+ 10 mM FBP), measuring fluorescence from excitation at both 405 nm and 488 nm (Figure 1C). After the first round of mutatagenesis and screening, the top four variants were used to initiate parallel evolution trajectories. Three rounds of error prone mutagenesis were completed on these parallel libraries, screening ∼10^3^ variants each round across the four libraries. For 8 variants per round (2 from each library), we benchmarked performance *in vitro* with a FBP titration (0-1 mM FBP), in mammalian cells with an abbreviated glycolytic stress test, or in cultured neurons with action potentials evoked by field stimulation. After the first four rounds of mutagenesis, the best performing variant from each library were pooled as templates for a StEP DNA shuffle reaction. In this step we identified an improved variant with a total of six mutations relative to HYlight, called HYlight2. HYlight2 exhibits improved intensity changes in the 488 channel, while retaining the ratiometric excitation and affinity properties of the original. This library also resulted in a series of improved sensors with shifted affinity, the best of which was retained and called HYlight_low_ (SI Figure).

**Figure 1.**
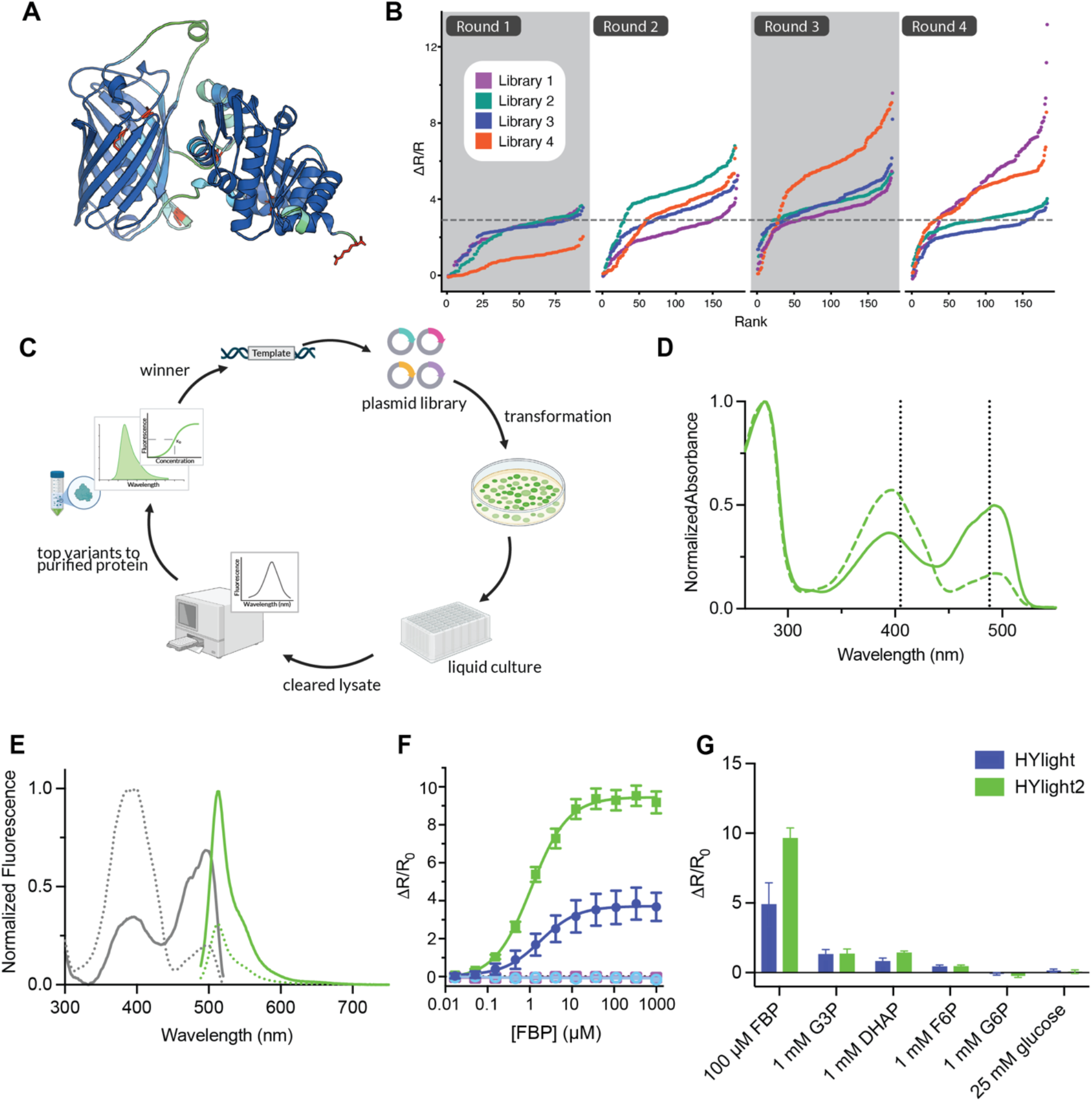
Overview of engineering and *in vitro* characterization of HYlight2. **(A)** AlphaFold3 model of HYLight2 colored according to pLDDT with changed resides identified from screening highlighted in red. **(B)** Library splitting of epPCR libaries allow each library to independently explore sequence space from a single starting scaffold. **(C)** Scheme of bacterial screening workflow. **(D)** Normalized absorbance spectra for HYLight2 with 6 mM FBP (solid line) or without FBP (dotted line). **(E)** Normalized excitation spectra (gray) with 6 mM FBP (solid) or without FBP (dotted) also plotted is normalized emission spectra (green) with excitation at 480 nm. **(F)** FBP titration curves of HYLight2 (green) and HYLight (purple) with their corresponding binding-dead versions (light blue and dark pink, respectively). **(G)** Selectivity for FBP over other glycolytic intermediates.

### In Vitro Characterization

We characterized the physicochemical properties of the sensors in purified protein to measure the binding affinity and molecular brightness. HYlight variants exhibit two absorption maxima at 396 nm and 492 nm corresponding to the protonated and deprotonated chromophore, respectively (Figure 1D, Table 1). Excitation into either band yields fluorescence with an emission maximum of 513 nm. HYlight functions as an excitation ratiometric sensor; as FBP concentration increases, the 490 nm band increases while the 400 nm band decreases (Figure 1E). We measured the quantum yield in the presence of 6 mM FBP at 490 nm finding that HYlight2 has a slightly lower quantum yield than HYlight (0.41 vs 0.51) (Table 1). However, the molar extinction coefficient of the saturated sensor at 490 nm was slightly higher than that of HYlight (67000 M^-1^cm^-1^ vs 53000 M^-1^cm^-1^) (Table 1). The apparent dissociation constant of HYlight2 for FBP is comparable to HYlight1 (1.10 ± 0.19 µM vs 2.16 ± 1.17 µM) (Figure 1F). However, the low affinity variant, HYlight_low_ exhibits a slightly lower ΔR/R_0_ of 6 and an order of magnitude higher apparent dissociation constant (18.9 ± 0.95 µM) (Figure S1). Thus, this variant may allow for detection of changes in FBP in cells with higher resting FBP levels where HYlight2 is saturated.

**Table 1.**
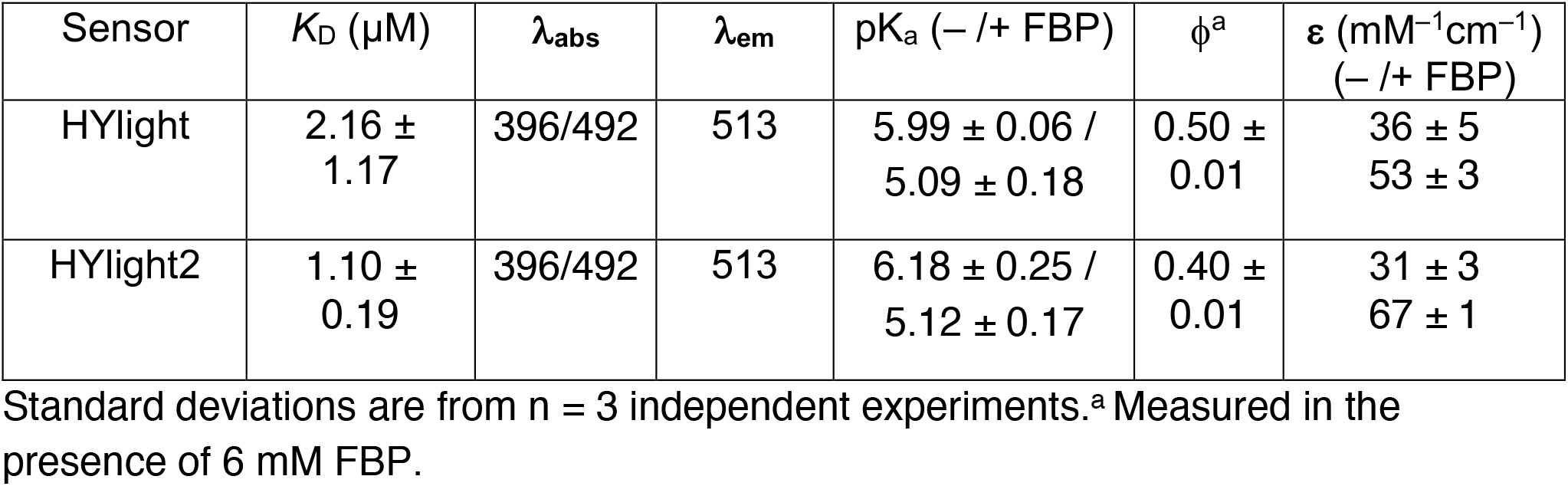
Photophysical parameters of HYlight1 and HYlight2.

HYlight2 displays a larger response to FBP with a ΔR/R_0_ of 9.5 ± 1.04 compared to 3.77 ± 1.43 for HYlight. It is worth noting that the fluorescence change (ΔF/F_0_) from excitation at 490 nm is substantial, so HYlight2 can also be imaged as a single wavelength intensiometric sensor. We then examined the response of HYlight2 as a function of pH, measuring the fluorescence ratio of 488/405 with and without FBP over the pH range 3.8-8.6. The unbound forms of HYlight and HYlight2 display a p*K*_a_ of 5.99 and 6.18, while the bound forms display a p*K*_a_ of 5.09 and 5.12, respectively (Figure S2). While the unbound forms of both sensors display only slight changes in the 488/405 ratio as a function of pH, the bound form of HYlight2 shows a more pronounced change as well as a higher fluorescence ratio when bound. Despite this increased pH sensitivity, the pKa is still relatively low, and a lowering of pH due to high glycolytic activity would result in a lower 488/405 ratio rather than spurious turn-on activity.

Lastly, we confirmed HYlight2 retains its high specificity for FBP over other glycolytic metabolites: dihydroxyacetone phosphate (DHAP), glyceraldehyde 3-phosphate (G3P), fructose 6-phosphate (F6P), glucose 6-phosphate (G6P), and glucose. Similar to HYlight, HYlight2 and HYlight_low_ show minimal nonspecific changes in fluorescence ratio (Figure 1G, Figure S1).

### Live cell imaging with HYlight2

To test the sensitivity of HYlight2 and HYlight_low_ in mammalian cells, we mimicked the common Seahorse XF protocol for a glycolytic stress test by starving the cells of glucose for 1 hour prior to imaging and then adding 11 mM glucose followed by 2.5 µM oligomycin and 16.7 mM 2-deoxyglucose (2-DG). The levels of FBP were monitored by measuring the fluorescence at 405 nm and 488 nm and the ratio change was calculated for the time course (Figure 2A, B). During a typical stress test, the cells exhibit a transient spike in FBP concentration upon glucose addition, followed by a plateau. Addition of oligomycin induced another increase in FBP levels, which was then reversed by the addition of 2-DG. Compared to HYlight, HYlight2 showed increased sensitivity towards the addition of glucose and oligomycin while retaining similar kinetics of responses. Overall, HYlight2 showed a ΔR/R_0_ of 4.9 in response to the initial glucose stimulation, in comparison to 1.2 for HYlight and 3.0 for HYlight_low_ (Figure S3).

**Figure 2.**
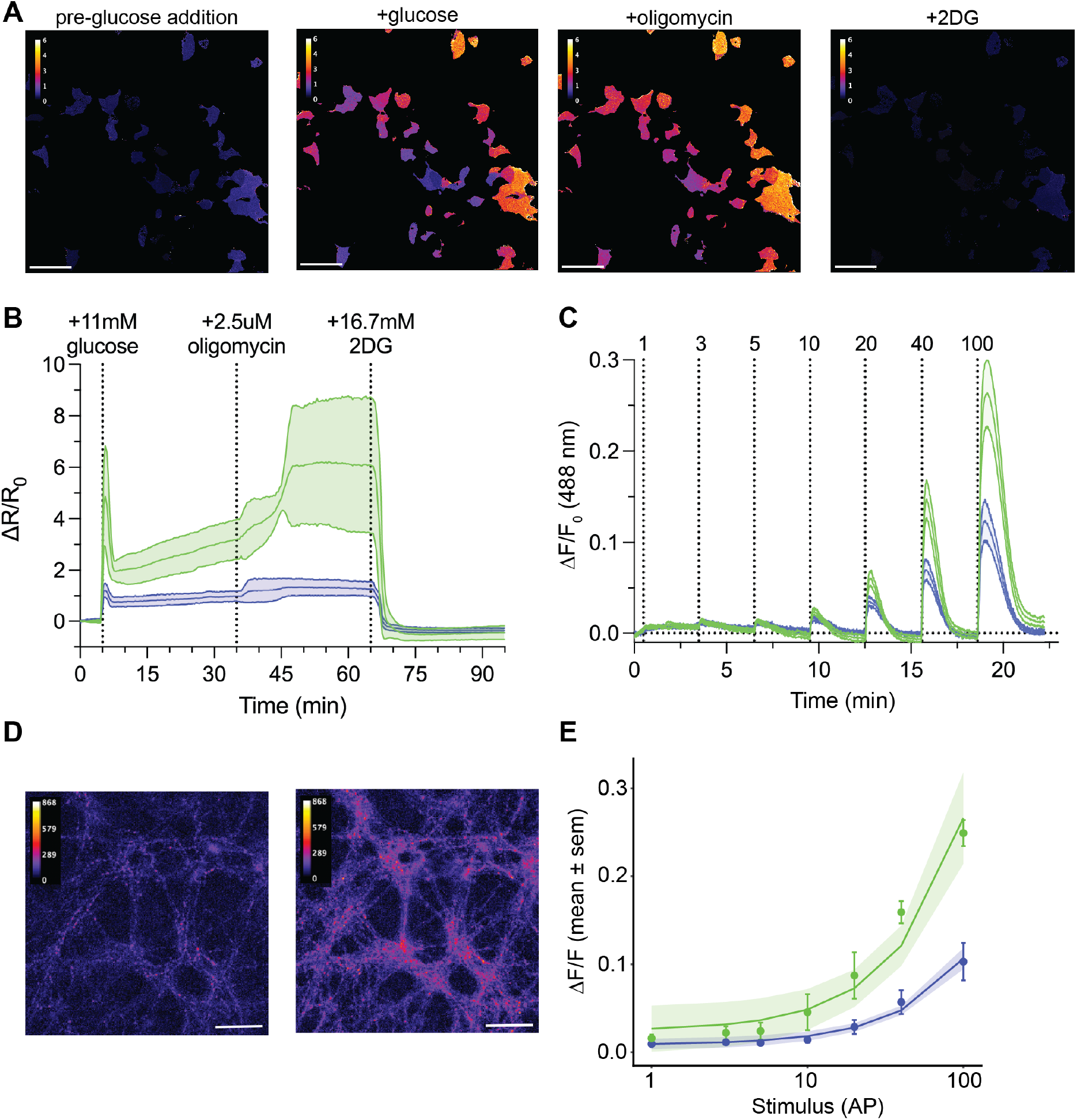
Glycolytic stress test and neuronal metabolic burden. **(A)** Representative images from a glycolytic stress test prior to glucose addition, after the addition of glucose, after oligomycin addition, and after 2-DG addition. **(B)** Plotted mean ΔR/R_0_ ± sem for glycolytic stress test from n = 6 replicates with 5-10 ROI per replicate. Green is HYlight2 and purple is HYlight. **(C)** Plotted mean ΔF/F_0_ ± SD from a representative sample of neurons (5 and 6 ROIs) stimulated with increasing numbers of action potentials. Green is HYlight2 and purple is HYlight. **(D)** Representative images from neurons with 0 AP (left) and 100 AP (right). **(E)** Linear regression with 95% CI on fit of sample means of ΔF/F_0_ vs stimulus for n = 2 experiments (10-12 ROIs). Green is HYlight2 and purple is HYlight.

**Figure 3.**
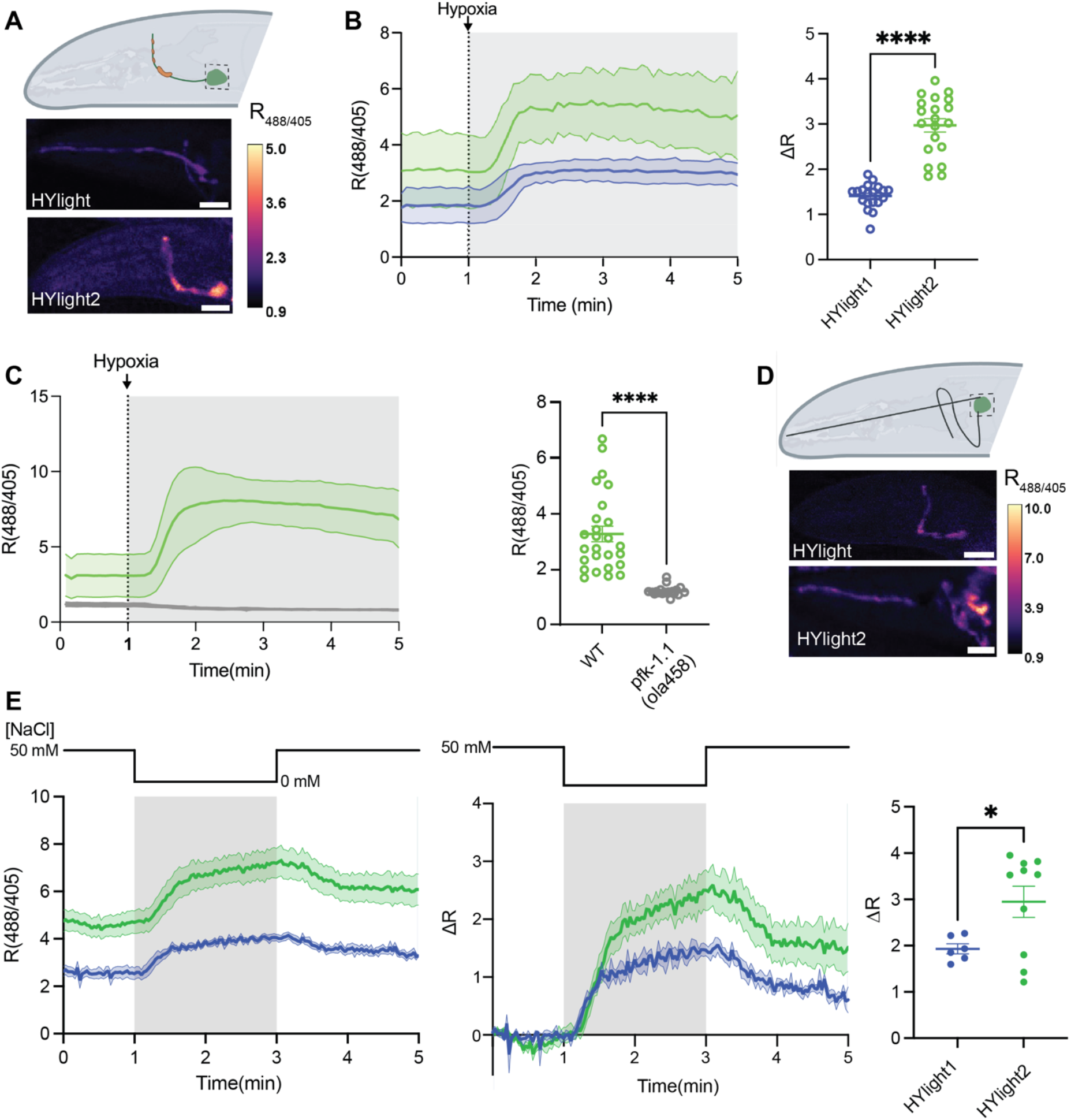
HYlight2 comparison in AIY and ASER neurons in *C. elegans*. **(A)** AIY interneurons respond to transient hypoxia by increasing glycolysis (top).^13^ Representative images of HYlight (middle) and HYlight2 (bottom) in AIY neurons, scale bars 20 µm. **(B)** Response of HYlight2 (green) and HYlight (purple) to transient hypoxia in AIY neurons over time (left). The change in ratio as a response to hypoxia in HYlight2 is significantly higher than that of HYlight (n=20 each, Mann Whitney U test, p<0.0001). **(C)** HYlight2 does not respond to hypoxia in a *pfk* knockout mutant (gray) (n=26 for HYlight1 and n=19 for HYlight2, Mann Whitney U test, p<0.001). **(D)** ASER neurons respond to changes in salt concentration.^12^ Representative images of HYlight (top) and HYlight2 (bottom) in ASER neurons, scale bars 20 µm. **(E)** HYlight2 exhibits higher ratios in ASER neurons as a response to lowering salt concentrations and exhibits a higher change in ratio than HYlight (n=6 for HYlight1 and n=10 for HYlight2, Welch’s t-test, p=0.0158).

### Imaging glycolytic flux after stimulated neuronal activity

Firing action potentials is metabolically costly, requiring neurons to rapidly replenish ATP used in neuronal terminals for the release of neurotransmitters at the synaptic cleft. The energetic demands of neuronal activity have been challenging to measure without sensitive biosensors that report on specific metabolic processes. We measured the response of HYlight2 targeted to synapses in response to increasing numbers of action potentials and compared that to the response of HYlight (Figure 2C). Over the entire range of action potentials, HYlight2 displayed notably higher ΔR/R_0_ and ΔF/F_0_.

Furthermore, a linear fit of the ΔF/F_0_ response of the biosensor to the number of action potentials evoked revealed that HYlight2 is 2.5 times more sensitive than HYlight for the detection of glycolysis at synapses (Figure 2D, Figure S4, Table S2).

### Imaging glycolysis in C. elegans

HYlight has previously been used *in vivo* in *C. elegans*, revealing different glycolytic states of neurons across the animal.^7^ To compare the sensitivity of HYlight2 to HYlight *in vivo* we expressed HYlight2 in AIY interneurons and induced hypoxia, which increases the glycolytic output of these neurons. As in other systems, HYlight2 displayed an improved sensitivity towards changes in glycolysis in comparison to HYlight. The ΔR was 0.67±1.21 and 1.84±2.12 respectively. To ensure the specificity of the response, we also expressed HYlight2 in the *pfk1*.*1(ola 458)* knock out, which removes the enzyme responsible for production of FBP. As expected, hypoxia did not lead to an increase in FBP and HYlight2 showed no response. This supports the conclusion that HYlight2 is specifically responding to FBP in these neurons and *in vivo*.

We then measured neuronal responses to a physiological stimulus. ASER neurons are sensory neurons that respond to lowered salt concentrations in the environment.^12^ We had previously observed increases in glycolysis upon increases of neuronal activity in ASER neurons.^713^ We expressed HYlight and HYlight2 in ASER neurons and measured the change in glycolysis triggered by lowered salt concentration. HYlight2 displayed a higher ratio at resting compared to HYlight and an increased change in ratio when the NaCl concentration was changed from 50 mM to 0 mM demonstrating that it is more sensitive to changes in neuronal glycolysis in response to physiological stimulation.

### Measuring changes in pancreatic beta islets in zebrafish

We next compared HYlight and HYlight2 performance in the pancreatic islet of larval zebrafish, another metabolically relevant context amenable to *in vivo* live imaging. Using the UAS-Gal4 promoter system, HYlight and HYlight2 were expressed by the pancreas-specific enhancer trap Gal4 line-Tg:Et in the primary islet of larval zebrafish (Figure 4B, B’, B”). As is expected from Tol-based transgenesis in the injected F_0_ generation, copy number variations of HYlight insertions produced intensity differences between cells within a single islet (Figure 4B’, B”, arrowheads). Simultaneous expression of mScarlet in the cytoplasm allowed us to normalize differences in brightness between cells using a per-cell normalization strategy. Furthermore, we verified that HYlight and HYlight2 expression was not limited to a specific cell type within the pancreas by fixing and staining larvae after live imaging with antibodies for glucagon and insulin hormones to label α cells and β cells in the pancreatic islet, respectively (Figure 4B’, B”).

**Figure 4.**
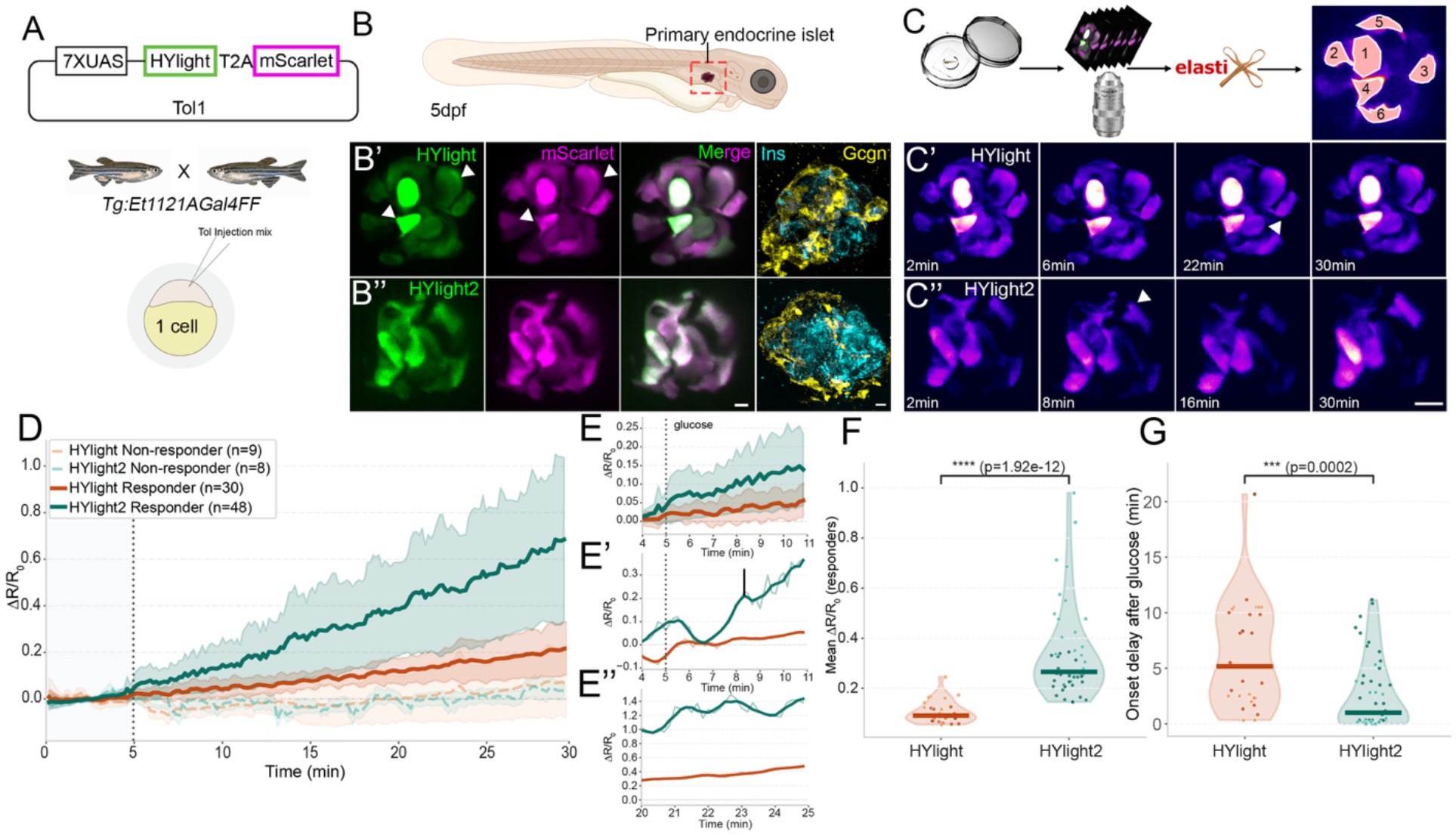
Characterization of HYlight2 responses *in vivo* in zebrafish pancreatic islets. (A) Plasmids and transgenic strategy used for injection of zebrafish embryos. **(B)** Imaging was done in the pancreatic islet of 5 dpf larval zebrafish. **(C)** Overview of the image processing pipeline. **(D)** Change in ΔR/R_0_ over time after glucose addition (dotted lines) for HYlight2 (green) and HYlight1 (red). Non-responding cells are plotted with dashed lines. **(E)** Inset of time course from 4 to 11 minutes for all cells and **(E’)** top responders, raw traces (thin line), traces smoothed using Savitzky-Golay filter (polynomial order = 2, window = 90sec). **(E”)** Inset of top responders at the end of the time course, raw traces (thin line), traces smoothed using Savitzky-Golay filter (bold line, polynomial order=2, window=90sec). **(F)** HYlight2 shows a higher mean ΔR/R_0_ (Mann Whitney U test, p = 1.92 e-12). **(G)** HYlight2 shows a shorter onset delay compared to HYlight1 (Mann Whitney U test, p = 0.0002).

To assess the sensitivity of each sensor, we performed *in vivo* recording of the islet during larval development with the addition of glucose. Volumetric imaging of HYlight and HYlight2 qualitatively revealed relatively slower dynamics of HYlight sensor when compared to HYlight2 (Figure 4C’, C”, arrowheads). After filtering out cells with high baseline noise (see Methods), cells pooled from all the recording sessions were categorized based on their response to glucose as responders or non-responders. Plotting the mean traces of non-responder and responder cells across the entire recording revealed that non-responders show a minimal or no response to glucose, with amplitudes fluctuating around 0.0 (Figure 4D, dashed orange and teal lines). HYlight responder cells reach a maximum amplitude of 0.2 with fluctuations plateauing by the end of the recording (Figure 4D, solid orange line). HYlight2, in contrast, ramped up to an average amplitude of ∼0.7 and continued to rise at the end of the recording (Figure 4D, solid teal line). HYlight2 displayed significantly larger continuous response amplitudes than HYlight across the length of the recording (Figure 4F, Mann Whitney U test, p=1.92e-12). Within shorter windows during recordings, the mean amplitudes of HYlight2 were higher than HYlight (Figure 4E), with the difference continuing to increase throughout the recording. In fact, the amplitude of the strongest responder cell of HYlight was 3 times lower than the strongest responder cell of HYlight2, both at the beginning (Figure 4E’) as well as at the end of the recording (Figure 4E”). Taken together, this indicates that the HYlight2 sensor has improved sensitivity for detection of changes in FBP compared to HYlight.

We estimated the onset delay kinetics for HYlight and HYlight2 to compare response rate of the two sensors (see Methods). Interestingly, HYlight2 sensed persistent FBP levels ∼4 times faster than HYlight as seen in Figure 4G (Mann Whitney U test, p=0.0002). This further demonstrates that HYlight2 has higher sensitivity towards repeated FBP fluctuations compared to HYlight.

### Characterization of glycolytic heterogeneity in mouse liver

To evaluate the performance of HYlight2 for *in vivo* detection of glycolytic activity, we performed intravital imaging of the liver, which exhibits a well-characterized metabolic gradient along the portal vein (PV) to central vein (CV) axis (Figure 5A). Glycolytic activity is enriched toward the pericentral region, providing a physiologically relevant benchmark for sensor performance^1^. AAV-mediated expression of both HYlight and HYlight2 in hepatocytes was confirmed by GFP and miRFP fluorescence (Figure 5B). Quantification of the average sensor-to-reference ratio at the single-cell level revealed a significant shift in the distribution of ratios between the two sensors (two-sided Mann– Whitney U test, p = 7.69 × 10^−3^) (Figure 5B and C). While the majority of hepatocytes exhibited ratios between 0.5 and 1.5, cells with more extreme ratio values were predominantly detected by HYlight2 (Figure 5D). To further investigate this difference, we compared the dynamic range of the two sensors. HYlight2 captured both lower and higher ratio values (0.23–3.68) compared to HYlight (0.31–1.64) (Figure 5E). Specifically, the lower bound (5th percentile) was reduced in HYlight2, while the upper bound (95th percentile and maximum values) was substantially increased, indicating improved sensitivity at both ends of the ratio distribution. Finally, spatial analysis of single-cell ratios as a function of distance from the CV revealed a clearer and more extended gradient in HYlight2 (Figure 5F). While both sensors detected elevated glycolytic activity near the CV with a gradual decrease toward the periportal region, HYlight2 exhibited greater heterogeneity and enhanced separation across the spatial axis. Together, these results demonstrate that HYlight2 provides an expanded dynamic range and improved sensitivity for detecting glycolytic activity *in vivo*, enabling more accurate resolution of metabolic heterogeneity across the liver lobule.

**Figure 5.**
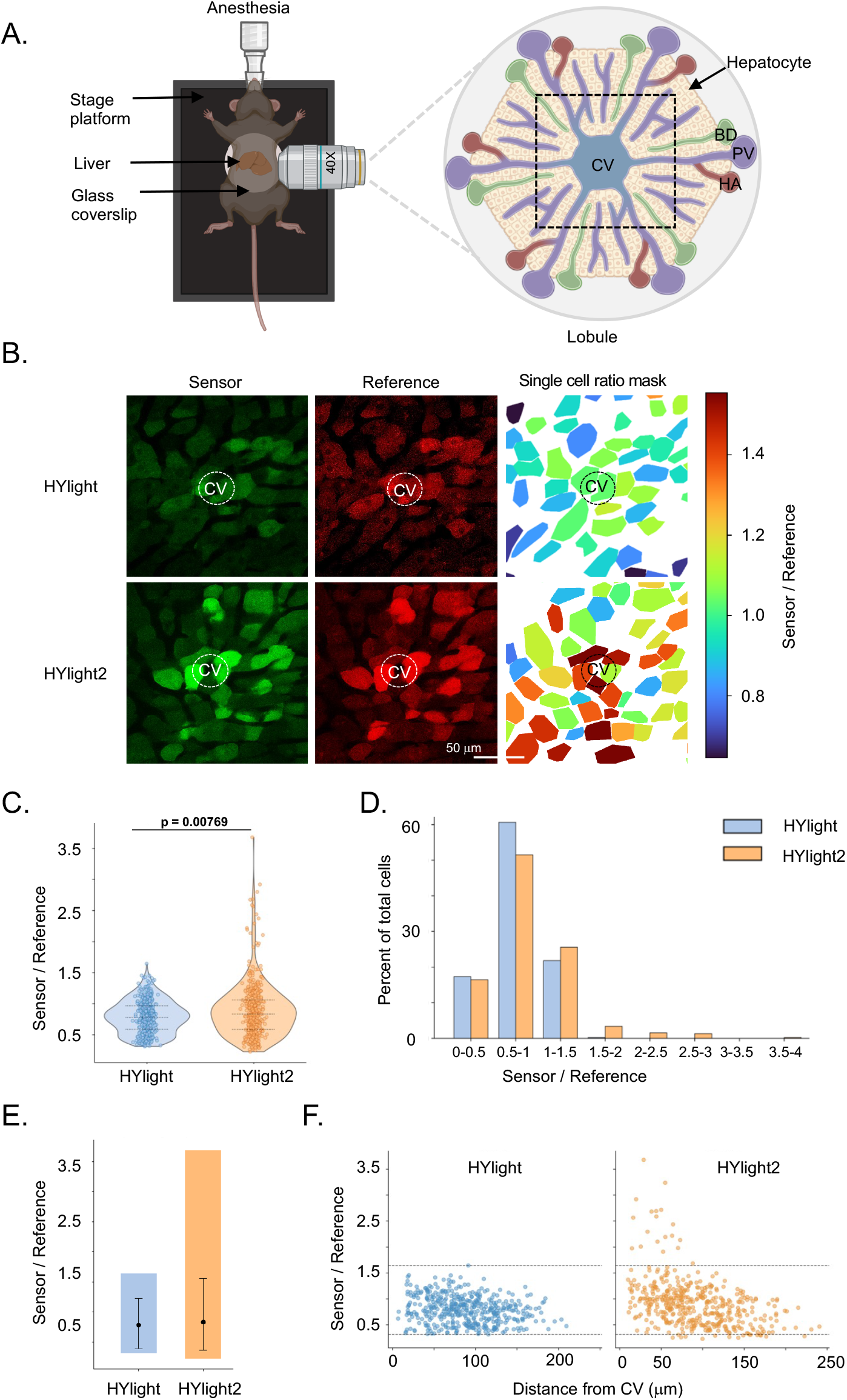
*In vivo* validation of HYlight2 in liver glycolytic gradient. **(A)** Schematic of the intravital imaging setup for monitoring glycolytic activity using HYlight sensors. The zoomed illustrates the hexagonal liver lobule, with portal triads consisting of the portal vein (PV), hepatic artery (HA), and bile duct (BD) located at the corners, and the central vein (CV) at the center. The imaging region near the CV is highlighted by a dotted box. The illustration was made using Biorender. **(B)** Representative fluorescence images of the sensor (GFP) and reference (miRFP) channels, along with corresponding single-cell ratio maps for HYlight1 and HYlight2. The position of the CV is indicated to visualize the spatial glycolytic gradient. The calibration bar on right indicates ratio values for corresponding color. **(C)** Violin plots showing the distribution of single-cell sensor-to-reference ratios. Individual cells are overlaid as points, and dotted lines indicate quartiles. HYlight2 data were randomly downsampled to match HYlight1 (n = 450 per group). Statistical significance was assessed using a two-sided Mann–Whitney U test (p = 7.69 × 10^−3^). **(D)** Clustered bar plots showing the percentage of cells within defined ratio bins. **(E)** Dynamic range comparison showing minimum, 5th–95th percentile, and maximum values for each condition. Black dot indicates the median value. **(F)** Scatter plot of single-cell sensor-to-reference ratios as a function of distance from the CV. Each point represents an individual hepatocyte. Dashed horizontal lines indicate the minimum and maximum values observed in HYlight1, providing a reference range for comparison.

## Discussion

This paper reports on an improved sensor for FBP, HYlight2, which can relay information about the glycolytic state of single cells. To engineer HYlight2, we explored sequence space using four separate directed evolution trajectories, eventually leveraging all the beneficial identified mutations in a recombination library. Each trajectory improved upon its initial sequence, but the trajectory producing the best final variants was not the one that led after the first round. This suggests that independent trajectories are a more efficient use of screening bandwidth and preserve diversity. In the end, six mutations were used to generate HYlight2. Notably, one of these mutations is adjacent to the FBP binding site (G92D) and two are proximal to the central alpha helix of the fluorescent protein (V263M, S267C). Two of these three mutations are present in both HYlight2 and HYlight_low_ supporting the idea that they contribute to the improved dynamic range of these sensors *in vitro*.

HYlight2 retains many of the favorable photophysical characteristics of HYlight, including ratiometric excitation and brightness adequate for *in vivo* imaging. Furthermore, our engineering efforts preserved affinity and selectivity for FBP, while identifying new affinity variants with useful dynamic range. This performance translated to mammalian cell culture, where a glycolytic stress test showed that HYlight2 outperforms HYlight in dynamic range. We also demonstrated the capability of the sensor to detect changes in glycolytic activity arising from the energetic burden of firing action potentials. By targeting the sensor to the synapse, it can report localized changes in glycolytic activity. In this setting, we found that HYlight2 is 2.5x more sensitive than HYlight, permitting the detection of glycolytic changes evoked by small numbers of action potentials. Similarly, HYlight2 improved the detection of altered glycolysis in *C. elegans* AIY and ASER neurons when the organisms increase glycolysis upon transient inhibition of oxidative phosphorylation (due to transient hypoxia) and upon physiological sensory stimulation, respectively. We expect this sensor will be broadly useful for understanding the energetic demands of neural activity, particularly at physiologically relevant levels of neural activity.

We also explored how this sensor can report on other physiological processes. We were able to express HYlight across multiple cell types in the pancreatic islet. After glucose addition, we observed that HYlight2 showed higher amplitude and faster response onset to this change in physiologic state. We examined its performance in pancreatic islets in developing zebrafish embryos. To our knowledge this is the first report of using a sensor for glycolysis *in vivo* in zebrafish pancreatic islets. Given the crucial role of the pancreas in sensing and regulating physiological state, we expect this sensor to be useful in understanding its activity *in vivo*.

The liver provides a well-established spatial gradient in metabolic activity along the portal vein (PV) to central vein (CV) axis, driven by directional blood flow and resulting gradients in oxygen, nutrients, and signaling factors.^14^ This zonated microenvironment gives rise to three distinct metabolic zones along the PV-CV axis, with glycolytic activity enriched in the pericentral region. Using this system as an *in vivo* benchmark, we find that HYlight2 exhibits a clear improvement over HYlight in capturing glycolytic activity at the single-cell level. While both sensors detect elevated glycolysis near the CV, HYlight2 reveals a broader distribution of ratios, including cells with both lower and higher values that are not well resolved by HYlight. This is reflected in its expanded dynamic range and increased detection of cells at the extremes of the distribution. Importantly, the enhanced performance of HYlight2 is consistent across both distribution-based and spatial analyses. The increased heterogeneity revealed by HYlight2 is consistent with recent reports of subzonal metabolic variability in hepatocytes.^15^ Organelle-based measurements in that work demonstrated that metabolic states vary within zones rather than between discrete zones — a pattern HYlight2 now resolves at the cellular level with a genetically encoded tool. Together, these results demonstrate that HYlight2 provides improved sensitivity and dynamic range for detecting glycolytic activity in vivo, enabling more accurate resolution of metabolic heterogeneity across the liver lobule.

We have demonstrated that HYlight2 is more sensitive to changes in glycolytic activity in cultured cells and neurons. Additionally, HYlight2 is useful *in vivo*, providing an avenue for dynamic measurements of glycolytic changes in metabolically important tissues such as the brain, liver, and pancreas. Many of these tissues have cell-level differences in glycolysis, such as neurons and astrocytes in the brain,^16^ and *in vivo* imaging with HYlight2 offers a route to disentangling the spatial and dynamic regulation of metabolism in these contexts. We anticipate that HYlight2 will enable a better understanding of how different cell types coordinate glycolysis to support tissue-level function across a broad range of organisms.

## Acknowledgements

We thank Helen Farrants and Tim Ryan for insightful discussion and input. We acknowledge the Media Prep facility, Primary Cell Culture, and Immortalized Cell Culture, Aquatics, and Vivarium facilities at Janelia Research Campus. The authors acknowledge the Howard Hughes Medical Institute for funding. This work was supported by National Institutes of Health grants to DC-R (R35NS132156).

## Data and Code availability

Plasmids have been submitted to Addgene and will be available with the following IDs: 255338, 255339, 255886, 255887. Data is included in the data supplement (in preparation). Data processing code is available at github.com/tebolab/hylight-paper.

