## Supplemental Information for "Improved sensors for fructose-1,6-bisphosphate enable in vivo imaging of glycolysis"

**Contents:**

**Supplemental Figures**

**Supplemental Tables**

**Methods**

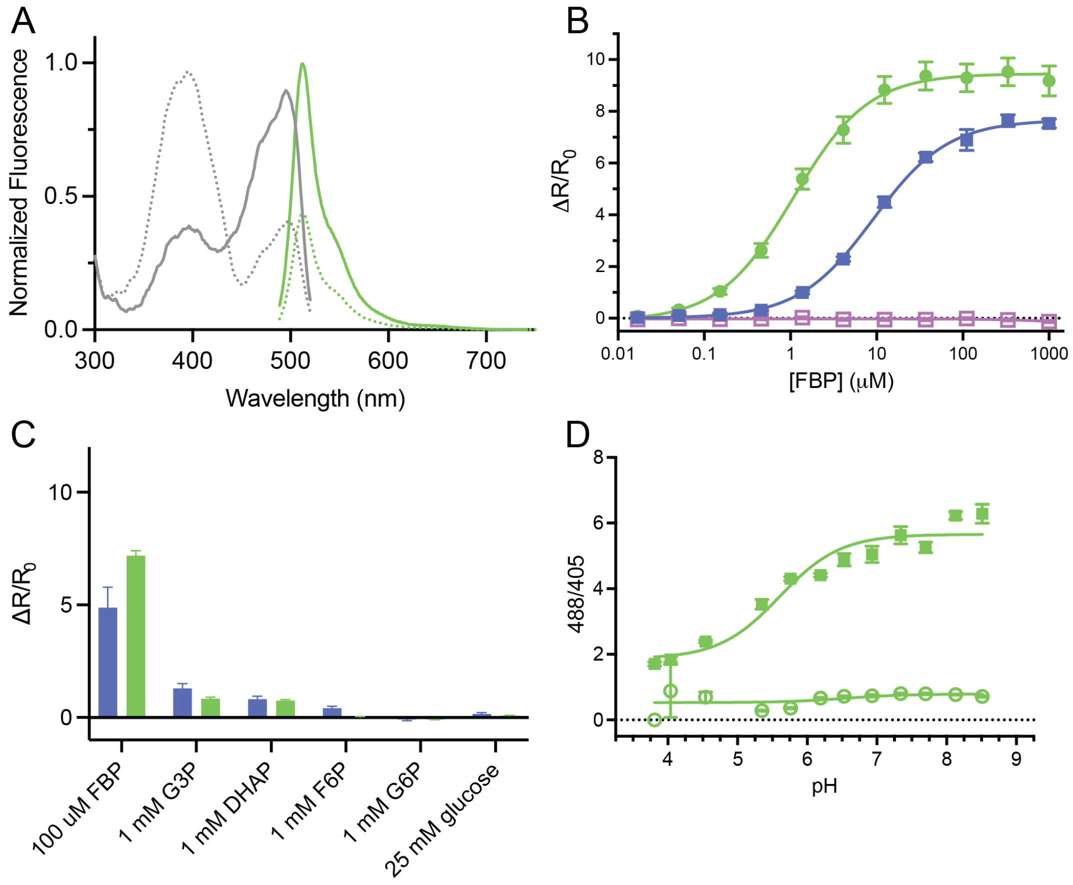


**Figure S1. In vitro characterization of HYlight$_{low}$. A)** Fluorescence excitation (gray) and emission (green) spectra of HYlight$_{low}$ with (solid line) and without (dotted line) FBP. **B)** FBP titration curves of HYight2 (same as Fig. 1F), HYlight$_{low}$ (purple), and HYlight$_{low}$-dead (dark pink). **C)** Selectivity of HYlight$_{low}$ for FBP over other glycolytic metabolites. **D)** p$K_a$ titration of HYlight$_{low}$ in the presence of FBP (solid circles) and without FBP (open circles).

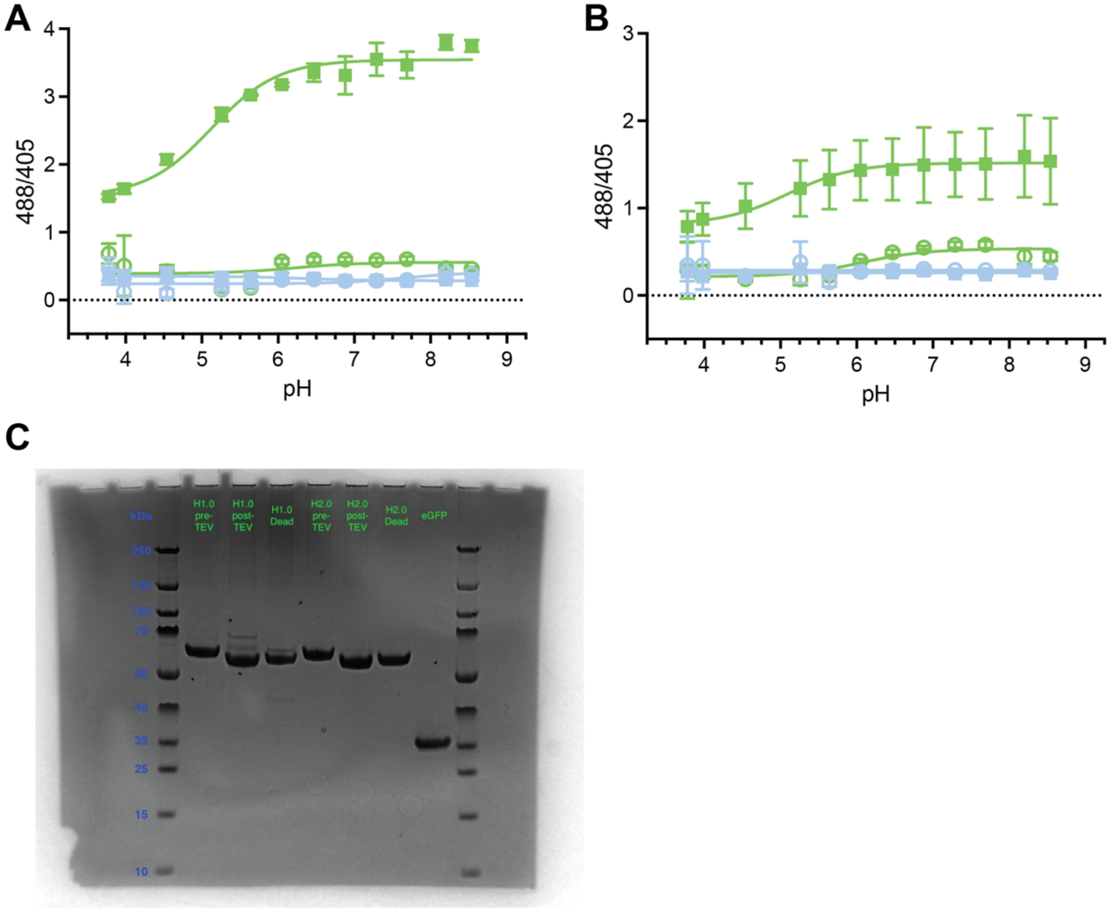


**Figure S2. Purification and p$K_a$ titrations for HYlight2. A)** p$K_a$ titration of HYlight2 in the presence of FBP (solid circles) and without FBP (open circles). **B)** p$K_a$ titration of HYlight1 in the presence of FBP (solid circles) and without FBP (open circles). **C)** SDS-PAGE of protein purification of HYlight1 and HYlight2.

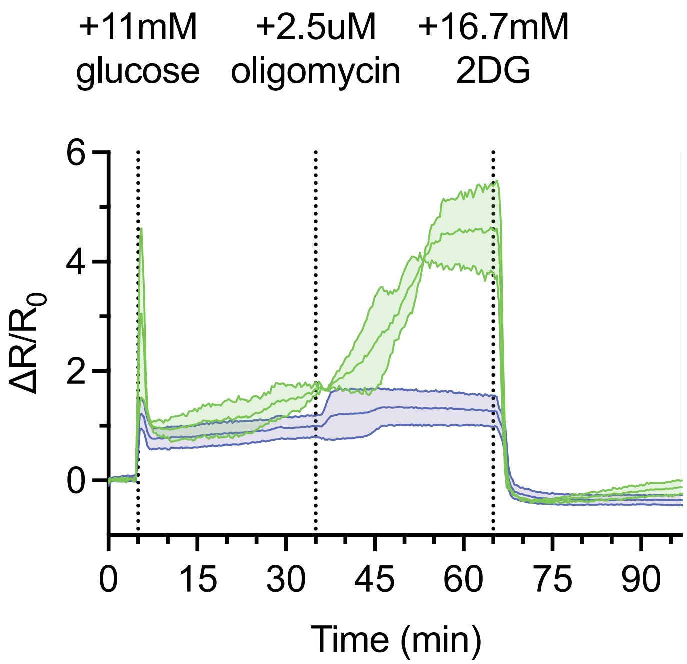


**Figure S3. Glycolytic stress test with HYlight$_{low}$.** Plotted mean $\Delta R/R_0$ ± sem for glycolytic stress test from n = 2 replicates with 5-10 ROI per replicate.

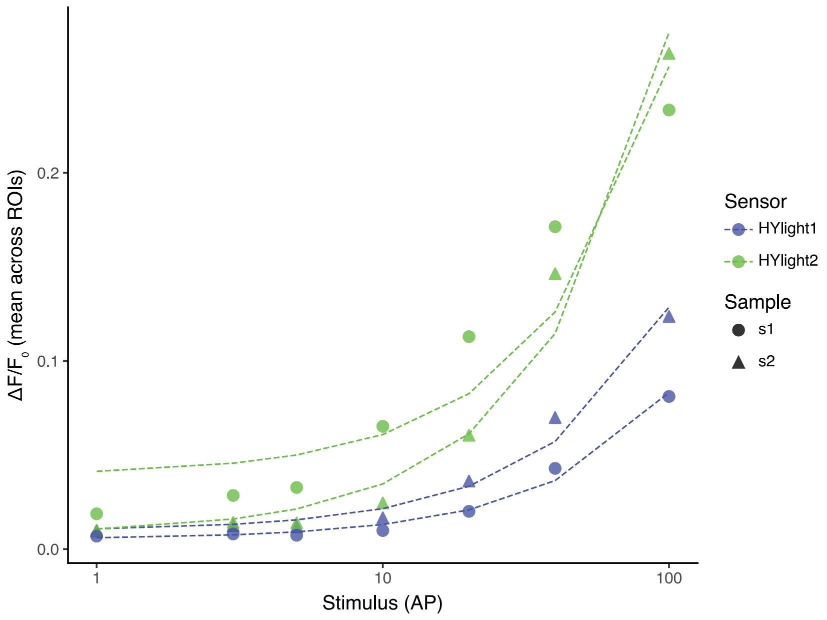


**Figure S4. Regression to individual replicates of neuronal stimulation data.** Linear regression fit (dotted lines) to individual replicates of neuronal stimulation for HYlight2 (green) and HYlight1 (purple).

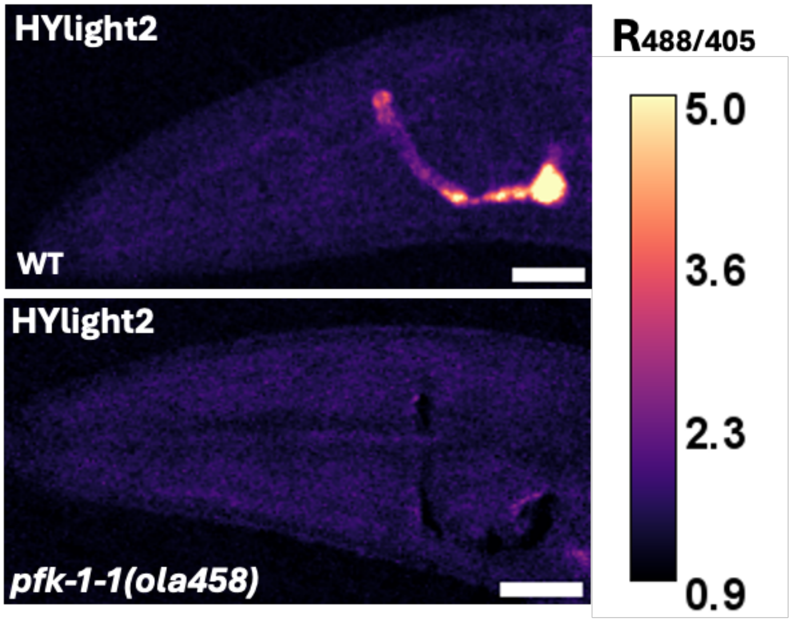


**Figure S5. Microscopy of HYlight2 in AIY neurons.** HYlight2 expressed in AIY neurons in a wildtype background (top) or pfk-1-1(ola458) background. Scale bars are 20 µm.

**Table S1. Photophysical parameters for $HYlight_{low}$**

| Sensor | $K_D$ (µM) | $\lambda_{abs}$ | $\lambda_{em}$ | $pK_a$ (– /+ FBP) | $\phi^a$ | $\varepsilon$ ($mM^{-1}cm^{-1}$) (– /+ FBP) |
|---|---|---|---|---|---|---|
| $HYlight_{low}$ | 2.16 ± 1.17 | 396/492 | 513 | 5.99 ± 0.06 / 5.09 ± 0.18 | 0.50 ± 0.01 | 36 ± 5<br>53 ± 3 |

Data are from n = 1 independent experiments with 3 technical replicates.[a] Measured in the presence of 6 mM FBP.

**Table S2. Summary of fit statistics for neuronal stimulation**

| Sensor | slope_mean | slope_sem | slope_n | t_stat | p_value |
|---|---|---|---|---|---|
| hylight1 | 0.000985 | 0.000205 | 2 | -4.471511 | 0.049359 |
| hylight2 | 0.002419 | 0.000246 | 2 | | |

**Methods:**

**Molecular Biology.**

Bacterial cloning: The open reading frame containing HYlight1.0 was PCR-amplified using Genemorph II random mutagenesis kit (Agilent), with a target mutation rate of 0-4.5 mutations/kb achieved by amplifying 500 ng DNA for 30 cycles of the brand suggested PCR program with a primer annealing temperature of 60ºC. Amplicons were run on an agar gel and purified with the Takara Bio Nucleospin Gel and PCR kit according to the brand protocol. The initial error prone library was inserted into a pET28aT7 vector via NEBuilder® HiFi DNA Assembly at 1:5 vector:insert. Then the libraries were expressed in chemically competent T7 Express E. coli cells (NEB) following the brand recommended transformation protocol onto LB agar plates with 50 uL/mL kanamycin. Sample plates were sent to GENEWIZ from Aztena Life Sciences for random colony Sanger sequencing to gauge mutational rate. Libraries are expressed on LB agar plates with 50 uL/mL kanamycin and 0.1 mM IPTG for screening. Based on lysate performance and sequence variation, a few variants are selected for large scale protein purification and in vitro testing. The best performing variants based on in vitro assays become parental sequences for separate, parallel error prone libraries. Each library undergoes the same cloning and screening procedure; about 200 variants per library (~800 total) were screened each round. Four rounds of error prone evolution were performed on four HYlight libraries in parallel. The final variant sequence from each library were input as template DNA for one round of staggered extension process PCR (StEP). About 200 ng total combined library DNA was amplified with NEB Taq 2X Master Mix in following PCR: initial denaturation 95ºC for 2min; 40 cycles of 95ºC for 20s, 59ºC for 30s, and 68ºC for 5s; extension at 68ºC then cool down at 10ºC, each for 2min; then 30 cycles of 95ºC for 20s, 62ºC for 30s, and 68ºC for 3min; and a final extension at 68ºC for 10min. The shuffled amplicon was restriction enzyme digested with NheI and XhoI and kit cleaned up. Insert and vector DNA were ligated with T4 DNA ligase (NEB) at room temperature for an hour. This DNA shuffled library was screened in lysate, and top performing variants were chosen for in vitro testing, one such variant being HYlight2.0.

Mammalian cloning: The open reading from containing HYlight variants was PCR-amplified using SeqAmp polymerase (Takara Bio). Amplicons were run on an agar gel and purified with the Takara Bio Nucleospin Gel and PCR kit according to the brand protocol. The plasmids were inserted into a CMV mammalian vector via NEBuilder® HiFi DNA Assembly at 1:5 vector:insert. Plasmids were expressed in chemically competent 5-alpha E. coli cells (NEB) following the brand recommended transformation protocol onto LB agar plates with 100 g/mL ampicillin. Bacterial colonies were picked and grown in 2 mL LB cultures for DNA clean up; resulting samples were sent for full plasmid sequence verification (Plasmidsaurus).

Synapse-localized HYlight variants: The open reading from containing HYlight variants was PCR-amplified using SeqAmp polymerase (Takara Bio). Initial amplicons were cleaned via PCR kit, then restriction enzyme digested with AgeI and NheI to match the vector. Digested amplicons were run on an agar gel and purified with the Takara Bio

Nucleospin Gel and PCR kit according to the brand protocol. The plasmids were ligated to an AAV viral vector with synpatophysin localization via T4 DNA ligase (NEB) at room temperature for 1 hour. Plasmids were expressed in chemically competent NEB Stable E. coli cells (NEB) following the brand recommended transformation protocol onto LB agar plates with 100 g/mL ampicillin. Bacterial colonies were picked and grown in 2 mL LB cultures for DNA clean up; resulting samples were sent for full plasmid sequence verification (Plasmidsaurus).

*C. elegans*: New HYlight variants identified from the in vitro screen were codon-optimized for expression in *C. elegans* using the GeneArt Gene Synthesis platform (Thermo Fisher Scientific). To further enhance expression, an intron was inserted within the first 100 bp of the codon-optimized sequence before synthesis. The resulting gene block was synthesised by and cloned into a *C. elegans* expression vector under the *pttx-3g* (Hobert et al., 1997, Neuron) and *flp6* promoters (Kim, K., & Li, C. (2004), Journal of Comparative Neurology), enabling expression of the HYlight 2 sensor in the AIY neuron and ASER neuron respectively. Plasmids were generated using Gibson Assembly (NEB).

Constructs were introduced into *C. elegans* by gonadal microinjection (Mello and Fire, 1995. For expression in the AIY neuron, the injection mix contained the ttx-3::HYlight2 expression plasmid (60 ng/µL), *pElt-7*::mCherry-NLS (15 ng/µL), and pSH5 empty vector (25 ng/µL). Stable transgenic lines were selected (ID: DCR10014) and used for subsequent experiments.

For expression in the ASER neuron, the injection mix contained flp6p::HYlight2 expression plasmid (60 ng/µL), gcy-5p::mCherry-NLS (15 ng/µL) and pSH5 empty vector (25 ng/µL). This generated a *C. elegans* strain (ID: DCR10023) stably expressing HYlight2 in ASE neurons. ASER was identified with the specific mCherry marker.

Zebrafish construct design: To generate plasmids with HYlight variants, 5XUAS_PACre_H2BmiRFP730 was used as the template for PCR reaction and HYlight was inserted using Gibson assembly. To express HYlight1 and HYlight2 in zebrafish pancreatic islet, 5XUAS-HYlight1-T2A-H2BmiRFP703 and 5XUAS-HYlight2-T2A-H2BmiRFP703 plasmids were used as a template to PCR amplify the Hylight1-T2A and HYlight2-T2A domains. These were combined into pTol1-7XUAS-mScarlet backbones using HiFi assembly to generate the 7XUAS-HYlight1-T2A-mScarlet and 7XUAS-HYlight2-T2A-mScarlet in Tol1 backbones.

**Error Prone Library Screening.**

The library was expressed in T7 cells on LB-agar plates containing 0.1 mM IPTG to induce protein production. About 400 colonies were visually picked under UV fluorescence into 96-deep well blocks (DWB) to inoculate 800 uL Terrific broth media. DWBs were left to shake 300 rpm at 37 °C until an OD/A600 of 0.6-8 was reached, approximately 4 hours. Cultures were induced with 0.5 mM isopropyl-β-D-thiogalactopyranoside (IPTG) then returned to shake at 300 rpm at 16ºC for 24 hours. Glycerol stocks were made using a small sample of culture and stored at -80ºC for future reference. Cells were pelleted in the DWBs by centrifugation at 4,000 × g for 10 minutes at 16 °C. The supernatant was discarded, and the bacterial pellets were resuspended with 0.5 mL 30 mM MOPS and 100 mM KCl, pH 7.5, then pelleted again for a total of three washes. The final pellet was lysed via freeze-thaw cycle between liquid nitrogen and a warm water bath, about 10 cycles. The lysed pellets are resuspended in 0.5 mL of buffer and centrifuged a final time. The resulting supernatant contains the protein, and aliquots are transferred from the screening DWB to a black chimney bottom 96-well plate. The plates are read on a BioTek Cytation 5 and excited at 488 nm and 405 nm with emission 525 nm while at ambient room temperature. Plates are initially measured, then dosed with 1 mM FBP from an auto-injector and re-read after 15 seconds of shaking to mix plate contents.

**Expression and purification of HYlight2 from *E. coli*.**

Chemically competent T7 Express *E. coli* cells (NEB) were thawed on ice and then transformed with plasmids encoding HYlight2.0 variants following the brand recommended transformation protocol. Transformed cells were plated onto LB-kanamycin agar plates and incubated overnight at 37°C. After incubation, one colony was picked and inoculated into 5 mL of LB-kanamycin media and then shaken at 225 rpm at 37°C overnight. The overnight culture was added to 500 mL of TB-kanamycin in a 2-liter flask. The large culture was shaken at 225 rpm at 37 °C until an OD/A600 of 0.6-8 was reached, approximately 3.5 hours. Induction with the addition of 0.5 mM isopropyl-β-D-thiogalactopyranoside (IPTG), then cultures were returned to shake at 225 rpm at 16ºC for 24 hours. Cells were pelleted by centrifugation at 4,000 × g for 20 minutes at 4 °C. The supernatant was discarded, and the bacterial pellet was kept on ice. Each sample was resuspended with 10 mL 30 mM MOPS and 100 mM KCl, pH 7.5, and transferred to a 50 mL conical tube. While on ice, the pellet was lysed by sonication for 5s on, 10s off, 3 minutes total, 70% amplitude. The lysate was transferred to high-speed centrifuge tubes and centrifuged at 30,000 × g for 60 min at 4 °C. The resulting supernatant, containing the clarified cell lysate, was collected in a 50 mL conical tube. Nickel-NTA resin for purification was prepared by taking 2 mL of resin slurry per sample and washing 3 times with MOPS buffer. The resin was resuspended in buffer and aliquoted such that each sample received 1 mL of packed resin to bind, and equal volume of 20 mM imidazole in MOPS buffer added then left for 1 hour, rotating at 4ºC. Samples were loaded into a Poly-Prep column and allowed the resin to settle. Two washes of ten column volumes 20 mM imidazole buffer were added, then His-bound protein eluted off resin with five column volumes of 500 mM imidazole in MOPS buffer and caught in Eppendorf tubes. Protein containing fractions were pooled and buffer exchanged back to MOPS buffer with PD-10 desalting columns. For tag-less preparation, protein samples were digested with TEV protease (Sigma-Aldrich) at a ratio of 1:50 TEV:protein by weight and dialyzed in a SLIDE-A-LYZER™ G3 dialysis cassette (Thermo Scientific) in 50 mM Tris-HCl, 1 mM EDTA, and 1 mM DTT, pH8.0, at 4ºC overnight. Nickel-NTA resin was added to the dialyzed proteins to bind un-cleaved protein and left for an hour while rotating at 4°C. The suspension was loaded onto a fresh Poly-Prep column, and the flow-through, containing tag-less HYlight, was collected. Proteins were concentrated in Amicon Ultra-15 Centrifugal Filter Unit and ~0.6 mL fractions were filtered in Costar Spin-X Centrifuge Tube Filters. Filtered protein fractions were run through a Superdex 200 Increase 10/300 GL column using a size exclusion protocol on an AKTA Avant FPLC machine. Selected fractions were pooled and concentrated to test in vitro.

**In Vitro Characterization.**

Apparent affinity: Proteins tested at 200 nM. Stock 1.02 mM FBP (octahydrate) solution in 30 mM MOPS and 100 mM KCl, pH 7.5, was serially diluted 1:3 to 17 nM, final FBP concentrations range from 1 mM to 16.9 nM and buffer only. Plated in black chimney 96-well plate, read on BioTek Cytation 5 and excited at 488 nm and 405 nm with emission 525 nm at room temperature.

pH: Proteins tested at 200 nM. Stock 102 mM FBP solution in 30 mM MOPS and 100 mM KCl, pH 7.5, diluted to 10.2 mM in various pH buffers (12 buffers ranging from pH 3.8 to 8.6) and tested final concentration of 10 mM FBP. Stock solutions to make pH buffers include citric acid and trisodium citrate for citrate buffers (pH 4.0–6.2), monosodium phosphate and disodium phosphate for phosphate buffers (pH 5.8–7.8), and tris-base and tris-HCl for tris buffers (pH 8–9.0). Proteins tested in pH buffers with or without FBP. Plated in black chimney 96-well plate, read on BioTek Cytation 5 and excited at 488 nm and 405 nm with emission 525 nm at room temperature.

Metabolite panel: Proteins tested at 200 nM. Following metabolites tested in 30 mM MOPS and 100 mM KCl, pH 7.5: 100 uM FBP, 1 mM G3P, 1 mM DHAP, 1 mM F6P, 1 mM G6P, 25 mM glucose. Plated in black chimney 96-well plate, read on BioTek Cytation 5 and excited at 488 nm and 405 nm with emission 525 nm at room temperature.

Spectra/QY: Proteins tested at 15 uM with or without 6 mM FBP solution, in 30 mM MOPS and 100 mM KCl, pH 7.5. Samples in 1 cm glass cuvette are diluted to absorbance at 490 nm of 0.04-0.08, then serially diluted 1:2 to acquire up to 5 data points of absorbance, excitation and emission spectra. Excitation spectra set at 600 nm, scanning 300-550 nm, and emission spectra set at 488 or 405 nm, scanning 550-750 nm. Data collected on Horiba Duetta Fluorescence and Absorbance Spectrometer at room temperature.

Extinction Coefficient: Proteins tested at 40 uM with or without 20 mM FBP solution, in 30 mM MOPS and 100 mM KCl, pH 7.5. Samples in 1 cm cuvette are diluted to absorbance at 490 nm of 0.2, measure with buffer or 1 M NaOH to denature protein folding. Data collected on Horiba Duetta Fluorescence and Absorbance Spectrometer at room temperature.

Photobleaching: Cover glass and coverslips were pretreated to make the surfaces hydrophobic by dipping them 2-3 times in Silane (Repel Silane, GE), followed by rinsing with ethanol and air drying them. To prepare the micro droplets; proteins (500 nM) was added in octanol by 1:9 ratio. The emulsions were mixed by hand tapping for 5 seconds. Further 5 uL of the droplet solution was sandwiched between a cover glass and a coverslip. Allowed to rest for few minutes so that the droplets are stable and not diffusing in the solution. One-photon bleaching experiments were performed with an inverted Nikon Eclipse Ti2 microscope. Images were acquired with 1 second of exposure every 2 seconds. Power at the objective was measured with a microscope slide power sensor to X mW, giving an irradiance of X mW mm−2. Each sample was bleached continuously for 10 minutes.

**Imaging and Quantification.**

<u>Mammalian Cell Culture:</u> HEK293WT cells were maintained in DMEM supplemented with 10% FBS and 1% L-glutamine. Cells passaged regularly upon growing to ~90% confluency until a cell passage number over 20 was reached then fresh cells were thawed. FuGENE transfection reagents used as per the manufacturer's protocol.

<u>Seahorse Glycolytic Stress Assay</u>: Approximately 200,000 HEK293WT cells were seeded in 35 mm ibidi glass bottom dishes treated with poly-D-lysine. Cells were transfected with 1 ug DNA for HYlight1.0 or HYlight2.0, using FuGENE transfection reagents and left for 48 hours. One hour prior to imaging, cells were glucose starved in Dulbecco's Modified Eagle Medium (DMEM) without D-glucose, L-glutamine, phenol red, or sodium pyruvate. Before imaging, cells were transferred from no glucose media to buffer containing 10 mM HEPES, 145 mM NaCl, 5 mM KCl, 1.2 mM $MgCl_2$, 2.6 mM $CaCl_2$, pH 7.4. Live cells were imaged on a Leica Stellaris 8 confocal microscope with a 40× glycerol immersion objective using the Adaptive Focus Control system. Cells were excited at 488 nm and 405 nm with emission monitored from 480 to 650 nm while at ambient room temperature. One image was acquired every 20 seconds. After establishing a five-minute baseline, 11 mM glucose was added to the cells and left to image for 30 minutes, then 2.5 uM oligomycin A was added and left for 30 minutes, and finally 16.7 mM 2-deoxy-D-glucose was added and left to image for 30 minutes.

Images were processed and quantified using Fiji ImageJ software to manually segment individual cells and generate ROIs (5-10 cells per sample). Mean intensity values for both the 488 and 405 nm channels were measured for each ROI and used to calculate the excitation ratio ($R = F_{488}/F_{405}$). The change in excitation ratio was calculated and plotted in Prism 11 software as $\Delta R/R_0 = (R_t - R_0)/R_0$, where baseline fluorescence intensity was calculated from the averaged intensity of the first 15 frames ($R_0$ = first 5 minutes or baseline) of imaging.

<u>Stimulation of Neuronal Metabolic Burden</u>: Rat hippocampal tissue was transfected with 0.5 ug DNA for HYlight1.0 or HYlight2.0 and cultured for 2 weeks in a glass bottom 24-well plate. Once ready for imaging, the cultures were washed with NGHKCM-1 buffer, then left in buffer containing synaptic blockers (10 uM CNQX, 10 uM (R)-CPP, 10 uM gabazine, 1 mM (S)-MCPG). Live cells were imaged on a Nikon Eclipse Ti2 widefield microscope with a 20× air objective with GFP filter settings at ambient room temperature. After establishing a 30 second baseline, neurons were stimulated

with 100 action potentials at 10 Hz for 10 seconds, imaged every second for 3 minutes, and stimulation was repeated once.

Minimum Stimulation of Neuronal Metabolic Burden: Rat hippocampal tissue was transfected with 0.5 ug DNA for HYlight1.0 or HYlight2.0 and cultured for 2 weeks in a glass bottom 24-well plate. Once ready for imaging, the cultures were washed with NGHKCM-1 buffer, then left in buffer containing synaptic blockers (10 uM CNQX, 10 uM (R)-CPP, 10 uM gabazine, 1 mM (S)-MCPG). Live cells were imaged on a Nikon Eclipse Ti2 widefield microscope with a 20× air objective with GFP filter settings at ambient room temperature. After establishing a 30 second baseline, neurons were stimulated with 1, 3, 5, 10, 20, 40, and 100 action potentials, with 3 minutes rest between each stimulation, and imaged every second for a total 23 minutes. Field stimulation at 10 Hz with a pulse width of 1 ms.

Images were processed and quantified using Fiji ImageJ software to manually generate ROIs (5-10 per sample) consisting of 3-5 synaptic boutons each. Mean intensity values for both the 488 and 405 nm channels were measured for each ROI and used to calculate the excitation ratio ($R = F_{488}/F_{405}$). The change in excitation ratio was calculated and plotted in Prism 11 software as $\Delta R/R_0 = (R_t - R_0)/R_0$ or $\Delta F/F_0 = (F_t - F_0)/F_0$ of the 488 nm channel only, where baseline fluorescence intensity was calculated from the averaged intensity of the first 30 frames ($R_0$ or $F_0$ = first 30 seconds or baseline) of imaging.

To assess sensor sensitivity, ROI traces were converted to $\Delta F/F_0$ using the mean fluorescence of the pre-stimulus baseline. For each stimulus block, the local baseline was defined as the 20 frames immediately preceding stimulus onset. The peak response was calculated as the maximum of a 5-frame rolling average within the stimulus window. ROI-level metrics were averaged within each sample, treating the sample as the independent unit of replication. Per-sample linear regressions of stimulus intensity vs. $\Delta F/F_0$ were fit (scikit-learn), and slopes were compared between HYlight2 and HYlight1 using a Welch's t-test. For visualization, sample-level means at each stimulus level were fit with a linear regression and a pointwise 95% confidence interval on the regression line.

Maintenance and Imaging *C. elegans*:

*C. elegans Maintenance.* Worms were maintained on NGM/agar plates at 20ºC using *E.coli* OP50 as the food source. L4 staged animals were picked on the previous day and imaging experiments were carried out on day 1 adults.

AIY Response to Hypoxia. For hypoxia experiments, 3% agarose pads were prepared and placed on a PDMS microfluidic device (Jang et al., 2021, Biophys J). Day 1 adult worms were mounted on the pads in 3 µL of M9 buffer containing 10 mM levamisole as a paralytic agent and covered with a square glass coverslip. The device was then mounted on the microscope and connected to the gas supply. Hypoxia was induced by flowing nitrogen through the microfluidic device, as described previously.

ASER Response to Physiological Stimulation. For checking the ASER neuron's response to salt-evoked physiological stimulations, a PDMS based microfluidics device was used (D. R. Albrecht et al., 2011, Nature Methods). Day 1 adult worms were loaded onto the device and imaged at 4x while the buffer salt concentration was changed from 50 mM to 0 mM as described previously (Singh et al., 2025, PNAS).

Imaging. Live *C. elegans* imaging was performed on a Nikon Ti2 equipped with a CSU-W1 spinning disk confocal unit and a Hamamatsu Orca-Fusion BT CMOS camera, with images acquired at 16-bit pixel depth. Ratiometric imaging of the HYlight FBP biosensor was achieved by alternating excitation between the 488 nm and 405 nm lasers with a fixed 525 nm emission filter. AIY-HYlight imaging was performed at 4× magnification with laser powers of 48% (488 nm) and 4% (405 nm), and an exposure time of 100 ms per channel.

All image analysis was performed in Fiji. For each image, background subtraction was applied to both the 488 nm and 405 nm channels. A region of interest (ROI) was drawn around the AIY cell soma in each channel, and the mean pixel intensity within the ROI was used as the channel value. The ratiometric value for each cell was then calculated by dividing the 488 nm ROI value by the 405 nm ROI value.

Zebrafish Maintenance

Adult zebrafish (*Danio rerio)* were maintained as per standard zebrafish husbandry procedures on a 14-10 light-dark cycle. The *Tg: Et1121AGal4FF* transgenic line was obtained from RIKEN, being maintained as part of the Kawakami collection.[1] Synchronously fertilized embryos from group mating crosses were used for microinjection of HYlight constructs. Injected embryos were maintained in accordance with the guidelines and procedures approved by the Janelia Research Campus Institutional Animal Use and Care Committee (Protocol No-25-0278.03).

Construct delivery and imaging of zebrafish pancreatic islets:

7xUAS-HYlight1.0-T2A-mScarlet and 7xUAS-HYlight2.0-T2A-mScarlet plasmids at 25ng/ µL concentration each were mixed with Tol1 mRNA (25ng/µL) to generate two injection mixes. 0.5nL of each injection mix was microinjected into 100-200 embryos of *Tg: Et1121AGal4FF*[2] transgenic line at the one celled stage across multiple experimental days. Embryos were screened for HYlight and mScarlet expression in the pancreas and maintained in filtered system water containing N-Phenylthiourea (PTU, 1X) starting 24 hours post fertilization (hpf) to prevent pigment formation until imaging.

Sample preparation: At 5 days post fertilization (dpf), larvae positive for HYlight1/2 and mScarlet expression in the pancreas were paralyzed with alpha-bungarotoxin (Thermo Fisher-B1601-6 µl droplet of 1 mg/ml stock per larva) for 90 seconds. Treated larvae were repeatedly washed with filtered system water and allowed to acclimatize for 10-15min in filtered system water before proceeding for imaging. Single larvae with minimal swimming activity were then individually embedded in 0.8% low-melting-point agarose (Invitrogen-16520-100, prepared in filtered system water) on MatTek (P35G-1.5-14-C) glass-bottomed dishes. Special care was taken to orient larvae laterally, with the right side of their body facing the glass bottom of the dish thereby positioning the pancreatic islet closest to the objective. The mounted larvae were submerged in filtered system water for 10-15min prior to imaging.

Imaging: Live timelapse imaging was performed on the Nikon CSU-W1 SoRA spinning disk confocal microscope, using a 40× (LWD, 1.15NA) water immersion objective in the dual camera setting. A 512x512 region of interest (ROI) encompassing the endocrine islet was defined for simultaneous excitation with 488 nm and 594 nm lasers at 25% power and 200 ms camera exposure time. Optical z-sections spaced at 5 µm intervals were acquired to cover the entire volume of the endocrine islet (9-10 sections) leading up to a net frame rate of 0.3Hz for the whole recording duration. At the $5^{th}$ minute of acquisition, D-glucose (Sigma Aldrich-G7021) solution to a final concentration of 75mM (prepared in filtered system water) was added to the dish which allowed for comparison of glycolytic responses to glucose stimulus v/s baseline fluctuations.

Image Processing: Raw timelapse recordings were corrected for motion artefacts using the Elastix registration pipeline[3], ideal for non-rigid registration. The mScarlet channel from registered recordings was used as reference for manual segmentation of ROIs around individual endocrine cells on Fiji[4]. Using a custom-made macro in Fiji, mean intensity values of z-plane specific ROIs from the HYlight and mScarlet channels were extracted across the entire recording, into Microsoft Excel for each larva imaged.

To extract fluorescence responses in endocrine cells a ratiometric quantification strategy was used. Each ROI's HYlight mean intensity was divided by its respective mScarlet mean fluorescence intensity at every timepoint of the acquisition to generate the ratio $R_t$. Median of R values over the first 5min of the recording (baseline before glucose addition) was calculated for each ROI and defined as $R_0$ for that ROI. This was then input into the following formula to get $\Delta R/R_0$ at every timepoint of the recording-

$$\frac{\Delta R}{R_0} = \frac{R_t - R_0}{R_0}$$

This $\Delta R/R_0$ metric represents the fractional change in glycolytic flux relative to the pre-stimulus baseline, normalised to the mScarlet reference channel to account for any residual motion or focus drift not corrected by registration. All individual cell (ROI) $\Delta R/R_0$ values across multiple embryos were then pooled for further filtering.

To eliminate cells with high baseline noise before glucose stimulus, we first calculated the standard deviation (SD) of $\Delta R/R_0$ for the 5min baseline window. A version of Median absolute deviation (MAD) was calculated from the SD distribution across all cells and scaled by multiplying factor 1.4826. Scaled MAD was then used in the following formula to define a threshold of acceptable noise during the pre-glucose baseline duration of the recording-

$$Threshold = Median\ of\ SD\ values + 3 * scaledMAD$$

All cells having baseline SD > Threshold were eliminated from further analysis.

Once cells with high baseline noise were excluded from the pooled dataset, we focussed on identifying cells that show a meaningful response to glucose stimulation. For this, we calculated mean $\Delta R/R_0$ during the post glucose addition period (25min) of the recording for each cell and input it into the following logical equation-

$$IF\ mean\frac{\Delta R}{R_0} > 3 * baseline\ SD, THEN\ "Responder", ELSE\ "Non-responder"$$

Once responder and non-responder cells were identified, mean $\Delta R/R_0$ values from multiples cells were combined. For plotting data from cells across different recording sessions, the time axis was resampled into 10sec grid by linear interpolation of the respective timeframes. Amplitude values across the entire distribution were statistically compared by Mann-Whitney U test. For assessing sensor dynamics, response onset delay was estimated. For this, each timepoint $\Delta R/R_0$ value was first compared with baseline SD for that cell (5 minute, pre-glucose) across the entire recording. The onset delay time for each cell was defined as the first timepoint during the recording when-

$$\frac{\Delta R}{R_0} > 3 * \text{ SD of baseline, sustained for consecutive timepoints spanning total 1min}$$

Mean onset delay times for all cells from HYlight and HYlight2 embryos were then plotted as a violin plot and compared by Mann Whitney U test for statistical significance. All data was analyzed on Python using the NumPy, pandas, SciPy packages and graphs for visualization were plotted using seaborn and matplotlib.

Immunostaining of imaged larvae: After live timelapse recording, larvae were unembedded from agarose and immediately fixed in 4% PFA (in PBS) overnight at 4°C. PFA was washed off with 1xPBS followed by permeabilization in PBS-TritonX100 (0.5%). The skin was carefully dissected with a fine pair of forceps to expose the internal organs. Larvae were then permeabilized and blocked in 8%Normal Donkey serum prepared in PBS-TritonX100 (0.5%) containing 1%DMSO (Sigma Aldrich 276855) for 3 hours at room temperature with slow rocking. Blocking solution was replaced by primary antibody solution containing Chicken anti GFP antibody (Aves Labs: GFP-1010,1:500), Rat anti-insulin antibody (R&D Biosystem: MAB1417, 1:200) and Rabbit anti-glucagon (Cell Signal:2760, 1:200) for overnight incubation at 4°C. The primary antibodies were further detected by secondary antibodies- Donkey anti-rat alexa fluor 405 (Thermo Fisher A48268), Donkey anti-chicken alexa fluor 488 (Thermo Fisher A78948) and Donkey anti-rabbit Alexa fluor 647 (Thermo Fisher A31573). Embryos were mounted in EasyIndex (EI-500-1.52) and imaged on 63X (1.4NA, oil) objective on Leica Stellaris confocal microscope.

Intravital Imaging of Mouse Liver:

Ten-week-old male C57BL/6J mice (strain #000664) from Jackson were used in this study. Animals were housed under a 12-hour light/dark cycle with ad libitum access to food and water. Approximately $4 \times 10^{11}$ adeno-associated virus serotype 8 (AAV8) particles encoding either HYlight1 (AAV.TBG.HYlight1.miRFPnano3—synthesized by VectorBuilder) or HYlight2 (AAV.TBG.HYlight1.miRFPnano3) were administered to separate mice via retro-orbital injection. To enhance AAV uptake by hepatocytes, mice (n = 2 per sensor) were fasted overnight and subsequently refed for 2 hours prior to viral injection. Intravital imaging of the liver was performed 7–10 days after AAV administration to allow sufficient expression of the constructs. Imaging was conducted between 9:00 am ± 2 hours for both conditions. All animal procedures were carried out in accordance with NIH guidelines and were approved by the Institutional Animal Care and Use Committee (Protocol #25-0280) at Janelia Research Campus, Howard Hughes Medical Institute.

Imaging was focused on regions surrounding the central vein (CV) of the liver because of the known bias of AAV-mediated expression toward pericentral hepatocytes. At least eight fields of view across two animals per construct were acquired for analysis. Images were collected using a Leica Stellaris 8 confocal microscope equipped with a 40X water immersion objective, using a pinhole size of 2 Airy units, a zoom factor of 2, and an image resolution of 1024 × 1024 pixels.

Sensor (GFP) and reference (miRFP) fluorescence images were acquired and used to generate single-cell sensor-to-reference ratio maps. Hepatocytes were manually segmented using Napari based on the reference fluorescence channel. For each segmented cell, mean fluorescence intensities were extracted, and the GFP/miRFP ratio was calculated. To ensure unbiased comparison between conditions, datasets were randomly subsampled to equal cell counts prior to statistical analysis. Differences between groups were evaluated using a two-sided Mann–Whitney U test. Spatial analysis was performed by calculating the distance of each cell from a manually defined reference point corresponding to the central vein location, which was identified based on anatomical features and visualization of blood flow during imaging. Data processing and visualization were performed using Python (NumPy, Pandas, Matplotlib).

**Primer Sequences:**

| Name | Sequence (5’ - 3’) |
|---|---|
| pET28aT7 vector forward (AGT72) | TAATAGCTCGAGCACCACCACCACC |
| pET28aT7 vector reverse (AGT71) | catgccctgaaaatacaggttttcGCTAGC |
| pET28aT7 error prone insert forward 1 (AGT73) | GCTAGCgaaaacctgtattttcagggcatgggcagcaaggatgttttgggtttg |
| pET28aT7 error prone insert forward 2 (JT41) | GCTAGCgaaaacctgtattttcagggcatgggcagcaaggatgttttggAtttg |
| pET28aT7 error prone insert reverse 1 (AGT74) | GGTGGTGGTGGTGCTCGAGCTATTAgcctgattcatctctcaacaactttttggcag |

| | |
|---|---|
| pET28aT7 error prone insert reverse 2 (JT39) | GGTGGTGGTGGTGCTCGAGCTATTAgcctCatt catctctcaacaactttttggcag |
| pET28aT7 error prone insert reverse 3 (JT40) | GGTGGTGGTGGTGCTCGAGCTATTAgcGtgatt catctctcaacaactttttggcag |
| pET28aT7 error prone insert reverse 4 (JT47) | GGTGGTGGTGGTGCTCGAGCTATTAgcctgTtt catctctcaacaactttttggcag |
| pET28aT7 StEP insert forward (JT59) | CCGCGCGGCAGCCATATGGCTAGCgaaaacct gtattttcagggcatg |
| pET28aT7 StEP insert reverse (JT60) | GTGGTGGTGGTGGTGCTCGAGCTATTA |
| pET28aT7 dead mutant forward (HFJpr1634) | gttgctgttactggtggtactgagattgaagctgttgct |
| pET28aT7 dead mutant reverse (HFJpr1633) | agcaacagcttcaatctcagtaccaccagtaacagcaac |
| pCMV insert forward (JT42) | TCCACTCGACACACCCGCCAGCGGCCGCG CCACCatgggcagcaaggatgtt |
| pCMV insert reverse (JT49) | TCAAGCTAGCAGAGGCCTGCGGATCCTTAgc Gtgattcatctctcaacaa |
| pAAV insert forward (JT64) | aatcaggatccaccggtcgccaccatggtcgactcatcaggca gcaaggatgttttg |
| pAAV insert reverse (JT69) | gatactttatacgaagttatgctagcTTAgcGtgattcatctctca acaa |

**List of Strains:**

| **Strain** | **Genotype** | **Source** |
|---|---|---|
| N2 | Bristol wild-type strain | CGC |
| DCR9089 | *olaIs138 [ttx-3p::HYlight1]* | Wolfe et al., 2024, PNAS |
| DCR9288 | *olais141[flp-6p::HYlight1]* | Singh et al., 2025, PNAS |

| | | |
|---|---|---|
| DCR10014 | *olaex5727[ttx-3p::HYlight2; elt-7p::mCherry-NLS]* | This study |
| DCR10023 | *olaex5729[flp-p::HYlight2; gcy-5p::mCherry-NLS]* | This study |
| DCR10024 | *olaEx5727; pfk-1.1(ola458)* | This study |

**HYlight2 Sequences.**

Sequence Color Legend:

CggR 96-180

Linkers

cpGFP

CggR 181-340

Mutations relative to HYlight1

HYlight2 contains the following amino acid changes: G92D, G125V, V263M, S267C, E406D, G505R

Protein Sequence:

MGSKDVLGLTLLEKTLKERLNLKDAIIVSGDSDQSPWVKKEMGRAAVACMKKRFSGKN
IVAVTGGTTIEAVAEMMTPDSKNRELLFVPARGDLGEPPYNVFIMADKQKNGIKANFKIR
HNIEDGVVQLAYHYQQNTPIGDGPVLLPDNHYLSVQSKLSKDPNEKRDHMVLLEFVTA
AGITLGMDELYKGGTGGSMVSKGEELFTGVVPILVELDGDVNGHKFSVSGEGEGDATY
GKLTLKFICTTGKLPVPWPTLVTTLTYGVQCFCRYPDHMKQHDFFKSAMPEGYIQERTI
FFKDDGNYKTRAEVKFEGDTLVNRIELKGIDFKEDGNILGHKLEYNFNKEDVKNQANTI
CAHMAEKASGTYRLLFVPGQLSQGAYSSIIEEPSVKEVLNTIKSASMLVHGIGDAKTMA
QRRNTPLEDLKKIDDNDAVTEAFGYYFNADGEVVHKVHSVGMQLDDIDAIPDIIAVAGG
SSKAEAIEAYFKKPRNTVLVTDEGAAKKLLRDESR*

DNA Sequence:

ATGGGCAGCAAGGATGTTTTGGGTTTGACCTTGTTGGAAAAGACTTTGAAAGAAAG
ATTGAACTTGAAGGACGCCATCATCGTTTCTGGTGATTCTGATCAATCTCCATGGGT
CAAAAAAGAAATGGGTAGAGCTGCTGTTGCTTGCATGAAGAAAAGATTTTCTGGTAA
GAACATCGTTGCTGTTACTGGTGGTACTACTATTGAAGCTGTTGCTGAAATGATGAC
CCCAGATTCTAAGAACAGAGAATTGTTGTTTGTTCCAGCTAGAGGTGATTTGGGTGA
ACCGCCTTATAACGTCTTTATCATGGCCGACAAGCAGAAGAACGGCATCAAGGCGA
ACTTCAAGATCCGCCACAACATCGAGGACGGCGTCGTGCAGCTCGCCTATCACTAC
CAGCAGAACACCCCCATCGGCGACGGCCCCGTGCTGCTGCCCGACAACCACTAC
CTGAGCGTGCAGTCCAAACTGAGCAAAGACCCCAACGAGAAGCGCGATCACATGG
TCCTGCTGGAGTTCGTGACCGCCGCCGGGATCACTCTCGGCATGGACGAGCTGTA
CAAGGGCGGTACCGGAGGGAGCATGGTGAGCAAGGGCGAGGAGCTGTTCACCGG
GGTGGTGCCCATCCTGGTCGAGCTGGACGGCGACGTAAACGGCCACAAGTTCAG
CGTGTCCGGCGAGGGCGAGGGCGATGCCACCTACGGCAAGCTGACCCTGAAGTT

CATCTGCACCACCGGCAAGCTGCCCGTGCCCTGGCCCACCCTCGTGACCACCCTG
ACCTACGGCATGCAGTGCTTCTGCCGCTACCCCGACCACATGAAGCAGCACGACTT
CTTCAAGTCCGCCATGCCCGAAGGCTACATTCAGGAGCGCACCATCTTCTTCAAGG
ACGACGGCAACTATAAGACACGCGCTGAGGTTAAGTTCGAGGGCGACACTCTGGT
TAACCGCATCGAGCTGAAGGGCATCGACTTCAAGGAGGACGGCAACATCCTGGGA
CATAAGCTTGAATATAACTTCAACAAGGAGGATGTTAAGAATCAAGCTAACACCATTT
GCGCTCATATGGCTGAAAAAGCTTCAGGTACTTACAGATTGTTATTCGTCCCAGGTC
AATTGTCTCAAGGTGCTTACTCTTCCATTATCGAAGAACCATCTGTCAAAGAAGTCTT
GAACACCATCAAATCCGCTTCCATGTTGGTTCACGGTATTGGTGATGCTAAAACTAT
GGCTCAACGCAGAAACACCCCATTGGAAGATTTGAAAAAGATCGATGATAACGATG
CTGTTACCGAAGCTTTCGGTTACTACTTTAATGCTGATGGTGAAGTTGTTCACAAGG
TTCATTCAGTTGGTATGCAATTGGATGATATTGATGCCATCCCAGATATTATTGCAGTT
GCTGGTGGTTCTTCTAAGGCTGAAGCAATTGAAGCTTACTTCAAGAAGCCAAGAAA
CACTGTTTTGGTTACTGATGAAGGTGCTGCCAAAAAGTTGTTGAGAGATGAATCACG
CTAA

$HYlight_{low}$ sequence: G92D, G125V, V143M, S267C, S398T, T416I, K498R, S504T

<u>Protein Sequence:</u>

MGSKDVLGLTLLEKTLKERLNLKDAIIVSGDSDQSPWVKKEMGRAAVACMKKRFSGKN
IVAVTGGTTIEAVAEMMTPDSKNRELLFVPARGDLGEPPYNVFIMADKQKNGIKANFKIR
HNIEDGVVQLAYHYQQNTPIGDGPMLLPDNHYLSVQSKLSKDPNEKRDHMVLLEFVTA
AGITLGMDELYKGGTGGSMVSKGEELFTGVVPILVELDGDVNGHKFSVSGEGEGDATY
GKLTLKFICTTGKLPVPWPTLVTTLTYGVQCFCRYPDHMKQHDFFKSAMPEGYIQERTI
FFKDDGNYKTRAEVKFEGDTLVNRIELKGIDFKEDGNILGHKLEYNFNKEDVKNQANTI
CAHMAEKASGTYRLLFVPGQLSQGAYSSIIEEPSVKEVLNTIKSATMLVHGIGEAKTMA
QRRNIPLEDLKKIDDNDAVTEAFGYYFNADGEVVHKVHSVGMQLDDIDAIPDIIAVAGGS
SKAEAIEAYFKKPRNTVLVTDEGAAKRLLRDETG*

$HYlight_{dead}$ sequence: T67E

MGSKDVLGLTLLEKTLKERLNLKDAIIVSGDSDQSPWVKKEMGRAAVACMKKRFSGKN
IVAVTGGTEIEAVAEMMTPDSKNRELLFVPARGDLGEPPYNVFIMADKQKNGIKANFKIR
HNIEDGVVQLAYHYQQNTPIGDGPVLLPDNHYLSVQSKLSKDPNEKRDHMVLLEFVTA
AGITLGMDELYKGGTGGSMVSKGEELFTGVVPILVELDGDVNGHKFSVSGEGEGDATY
GKLTLKFICTTGKLPVPWPTLVTTLTYGVQCFCRYPDHMKQHDFFKSAMPEGYIQERTI
FFKDDGNYKTRAEVKFEGDTLVNRIELKGIDFKEDGNILGHKLEYNFNKEDVKNQANTI
CAHMAEKASGTYRLLFVPGQLSQGAYSSIIEEPSVKEVLNTIKSASMLVHGIGDAKTMA
QRRNTPLEDLKKIDDNDAVTEAFGYYFNADGEVVHKVHSVGMQLDDIDAIPDIIAVAGG
SSKAEAIEAYFKKPRNTVLVTDEGAAKKLLRDESR*

1. Asakawa K, and Kawakami K. The *Tol2*-mediated Gal4-UAS method for gene and enhancer trapping in zebrafish. (2009) DOI: http://doi.org/10.1016/j.ymeth.2009.01.004
2. Yang Y H C, Briant L J, Raab C A, Mullapudi S T, Maischein H-M, Kawakami K, and Stainier D Y. Innervation modulates the functional connectivity between pancreatic endocrine cells. (2022) DOI: http://doi.org/10.7554/eLife.64526
3. Klein S, Staring M, Murphy K, Viergever M A, and Pluim J P W. elastix: A Toolbox for Intensity-Based Medical Image Registration. (2010) DOI: http://doi.org/10.1109/TMI.2009.2035616
4. Schindelin J, Arganda-Carreras I, Frise E, Kaynig V, Longair M, Pietzsch T, Preibisch S, Rueden C, Saalfeld S, Schmid B, Tinevez J-Y, White D J, Hartenstein V, Eliceiri K, Tomancak P, and Cardona A. Fiji: an open-source platform for biological-image analysis. (2012) DOI: http://doi.org/10.1038/nmeth.2019
